# Warming-induced sex bias underlies rapid recovery but may generate demographic imbalance in sea turtles

**DOI:** 10.64898/2026.01.16.699440

**Authors:** Fitra Arya Dwi Nugraha, Rebekka Allgayer, James Gilbert, Laura Sivess, Kirsten Fairweather, Artur Lopes, Sandra M. Correia, Berta Renom, Albert Taxonera, Stephen J. Rossiter, Justin Travis, Christophe Eizaguirre

## Abstract

Global warming threatens species with temperature-dependent sex determination (TSD) by increasing the risk of extreme offspring sex-ratio bias. In sea turtles, warmer incubation conditions produce more females. Such biases can transiently enhance population growth, while potentially generating demographic imbalance if male scarcity intensifies. Here, we combined 16 years of field monitoring, drone-based sex-ratio surveys, satellite-relayed telemetry, and eco-physiological modelling to comprehensively test whether rising temperatures can underpin the rapid increase in loggerhead (*Caretta caretta*) nesting in Cabo Verde, one of the world’s largest populations. We then tested whether the temperature–demography relationship found in Cabo Verde holds in 28 other loggerhead populations worldwide. In Cabo Verde, historical air temperature, lagged by one generation, was the strongest predictor of current nest counts, consistent with past warming increasing female recruitment into the breeding population. No other environmental characteristics in the breeding or foraging grounds correlated with nest counts. Independently, drone surveys revealed an approximately 9:1 female-to-male breeding sex ratio across two consecutive breeding seasons, while simulations reproduced increasing nesting trajectories under historical warming. Across global loggerhead populations, temperature–demography relationships were widespread and context-dependent, varying with latitude and thermal regime. Overall, these combined lines of evidence reveal a potential paradox of climate-driven recovery in TSD species: warming can increase female recruitment and apparent population growth, while continued directional warming may progressively increase the risk of demographic imbalance through male scarcity.

## Introduction

Global climate change is reshaping population dynamics across ecosystems, driving species to shift their ranges, adapt or decline, and in some cases to extinction (Nogués-Bravo et al. 2018; Weiskopf et al. 2020). Identifying the mechanisms by which climate warming influences population trajectories is therefore critical for understanding whether apparent population increases represent persistent recovery or transient responses to environmental change.

Among the most climate-sensitive taxa are species with temperature-dependent sex determination (TSD), in which incubation temperature irreversibly determines offspring sex (Lockley and Eizaguirre 2021). In some TSD reptiles, including all sea turtle species, warmer temperatures increase the proportion of females within a nest (Bull and Vogt 1979; Yntema and Mrosovsky 1982). While this bias can transiently enhance population productivity because more females can contribute directly to reproduction, it risks long-term demographic instability when male scarcity limits fertilization success (Fuentes et al. 2024; Laloë et al. 2014, 2024). Locally, extreme sex-ratio bias has been detected in Australian populations of the green turtle (*Chelonia mydas*), with over 90% of both juveniles and subadults being female (Jensen et al. 2018). However, not all populations seem to experience such extreme female bias (Pilcher et al. 2025). Seasonal and interannual variation in nest temperatures, among-clutch variation in thermal tolerance and local adaptation in pivotal temperatures may buffer warming-induced sex-ratio bias (Kynoch et al. 2024; Laloë et al. 2024; Santidrián Tomillo 2024; Tromp et al. 2025). The potential for both enhanced population growth and emerging demographic vulnerability makes TSD species informative models for understanding the demographic consequences of climate warming.

In sea turtles, temperature can influence not only hatchling sex ratios (Godfrey and Mrosovsky 2006) but also reproductive physiology (Fleming et al. 2020) and phenology (Mazaris et al. 2013; Nugraha et al. 2026), although phenological shifts are unlikely to keep pace with continued climate warming (Fuentes et al. 2024; Laloë and Hays 2023). Elevated sea surface temperatures (SST) can shorten inter-nesting intervals and enable more nesting events per season (Weber et al. 2011), while ocean productivity influences energy acquisition and thus reproductive frequency (Ramírez et al. 2021; Valverde-Cantillo et al. 2019). Consequently, increasing nest counts may reflect contemporary reproductive conditions, changes in survival or movement, conservation-driven recovery, or delayed recruitment of female-biased cohorts produced under warmer conditions (Hays et al. 2024; Mazaris et al. 2017).

The Cabo Verde archipelago hosts one of the world’s largest nesting aggregations of loggerhead turtles (*Caretta caretta*), where the number of nests has increased dramatically over the past decade (Roch et al. 2026; Taxonera et al. 2022). This rise has occurred alongside conservation efforts, including night patrols to deter poaching, initiated in the early 2000s and intensified around 2010 (Nugraha et al. 2026; Roch et al. 2026), and against a background of rapid regional warming since the late 1960s. Whether this recovery represents a demographically healthy rebound or is partly influenced by female-biased cohorts reaching maturity remains unknown. We hypothesise that conservation and contemporary ocean conditions alone cannot explain the magnitude and timing of the recent increase, which also reflects air temperatures at the time current breeders hatched. Warmer conditions may have produced more female hatchlings, resulting, decades later, in greater recruitment of females into the breeding population and consequently more nests.

We tested this hypothesis by integrating complementary datasets that address successive steps of a demographic pathway. Sixteen years of systematic nest monitoring on Sal Island (2008–2023) quantified the recent population increase in nest counts. Satellite-relayed telemetry from adult females tracked approximately 10 years apart was used to identify major post-nesting foraging areas and to assess whether their broad locations remained consistent through time before testing whether sea surface temperature and chlorophyll-a (of foraging and breeding grounds) correlated with nesting trends. Historical air-temperature data from the nesting grounds were then used to reconstruct thermal conditions experienced during the likely hatching period of current breeders, allowing us to test whether warming coincided with the recruitment of female-biased cohorts. We then developed an eco-physiological population model to test whether historical warming, through its effects on temperature-dependent sex determination and recruitment, could reproduce the observed increase in nesting activity. This model was contrasted against a null-model without the warming trends.

This structured hypothesis-driven approach generated a further demographic prediction: populations experiencing warming-driven female-biased recruitment should exhibit strongly female-biased breeding aggregations. Because males generally do not come ashore and so are missed by beach-based surveys, we used drone surveys over nearshore breeding aggregations to quantify the breeding sex ratio (Schofield et al. 2017). To determine whether this pattern represented a consistent demographic feature rather than a single-year anomaly, surveys were conducted across two consecutive breeding seasons (2022 and 2023). Finally, if temperature-dependent recruitment contributes more broadly to loggerhead population dynamics, similar relationships between historical temperature and subsequent nesting trends should be detectable beyond Cabo Verde. We therefore extended the analysis to 28 additional loggerhead populations worldwide to assess how temperature–demography relationships vary across geographic and thermal regimes (Figure 1A). Together, these complementary datasets allow us to distinguish contemporary environmental drivers of reproductive output from delayed demographic effects expected under temperature-dependent sex determination.

**Fig. 1:**
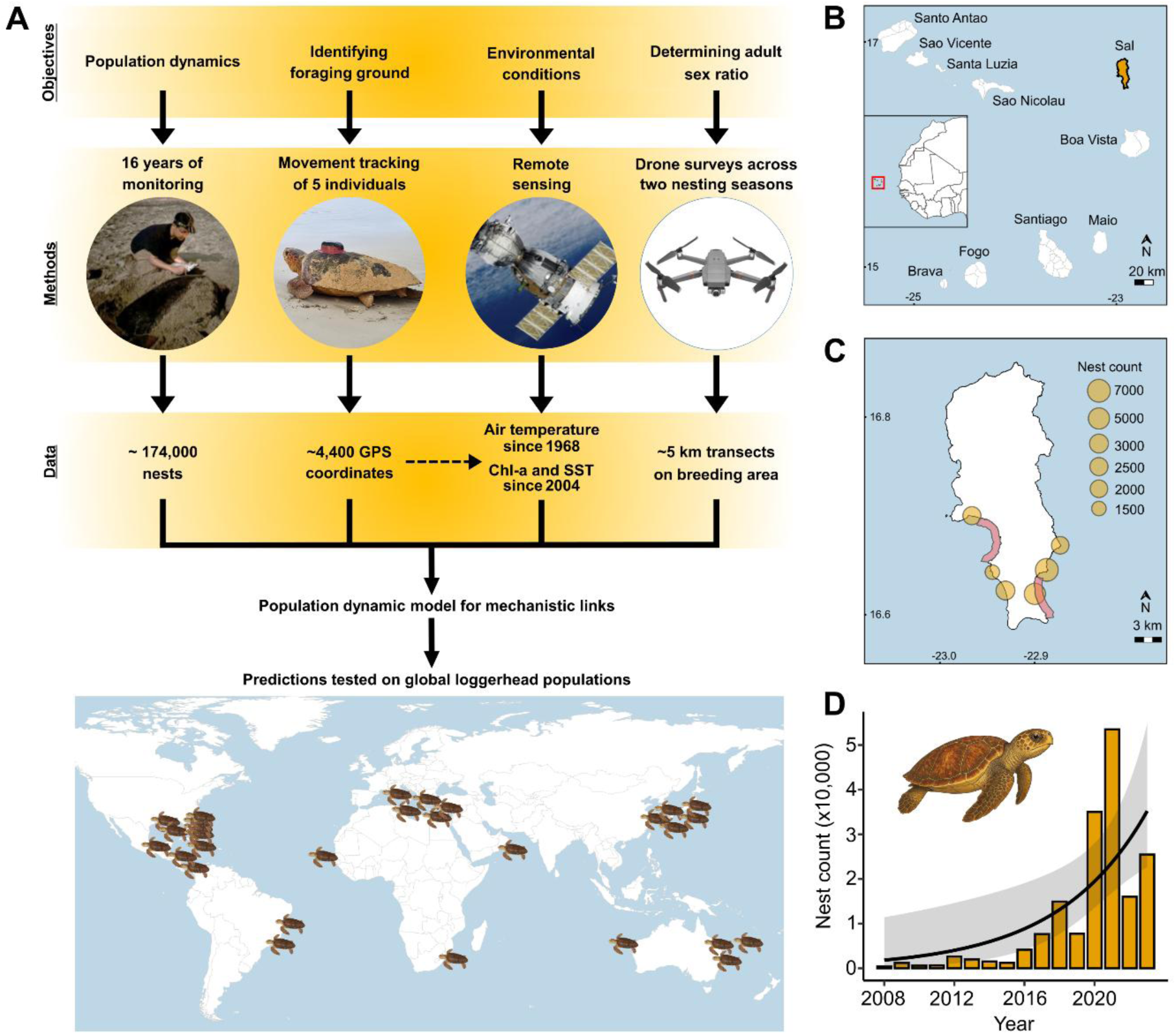
Study framework, study sites and loggerhead population trend. **A.** Study framework integrating multiple datasets to assess the ecological drivers of loggerhead population dynamics. Demographic changes were captured by long-term surveys; Satellite-relayed telemetry of nesting turtles was used to assess the broad stability of post-nesting foraging areas and to identify the regions from which environmental parameters were extracted; drone monitoring confirmed the breeding sex ratio over two consecutive breeding seasons. Broad predictions stemming from the relationships between population trends and historical air temperatures were tested across global loggerhead populations. Turtles on the map represent nesting populations included in this study (please refer to the page). **B.** *Main map*: the islands of the Republic of Cabo Verde, with Sal Island highlighted in yellow. *Inset map*: the Republic of Cabo Verde in relation to the African mainland (within the solid red square). **C.** Sal Island and main nesting sites with the size of the circle correlating with the average yearly nest counts from the period 2008-2023. Red areas near the coastline show where drone transects were conducted, Murdeira Bay in the west and Costa Fragata beach in the east. **D.** The positive trend is best supported by an exponential model (AICc= 347.86; β = 0.197 ± 0.071, t = 2.78, *p* = 0.015).

## Material and methods

### Nest Data Collection and Mark-Recapture Programme

Nesting season monitoring was conducted in the Republic of Cabo Verde, a volcanic archipelago composed of 10 islands and several islets, located ∼570 km off the coast of West Africa (Fig. 1B). Our study focused on the loggerhead nesting aggregation on Sal Island, which has been monitored systematically since 2008 (Fig. 1C). Foot patrols were conducted by members of SOS Tartarugas (2008–2014) and Associação Projeto Biodiversidade from 2015 onwards (https://www.projectbiodiversity.org) during the nesting seasons, from late June to October (Hays et al. 2022; Nugraha et al. 2026). This resulted in a dataset running from 2008 to 2023. The field monitoring covered approximately 25–27 km of nesting coastline, with variation due to the coverage of low-density nesting areas. During these night patrols, nests were counted and their GPS coordinates recorded daily. To avoid double counting, all nesting tracks were marked by drawing a large “S” across both the incoming and outgoing tracks.

Alongside beach patrols, an intensive mark–recapture programme has been conducted, in which individuals handled during the patrol were checked for the presence of a Passive Integrated Transponder (PIT) tag and, if untagged, fitted with a new one (Nugraha et al. 2026). This programme provided information on individual nesting behaviour and allowed the identification of turtles previously tagged at other nesting sites within Cabo Verde. Although loggerhead turtles generally show strong nesting-site fidelity (Baltazar-Soares et al. 2020), individuals may also use alternative nesting sites within or among reproductive seasons (Hays and Sutherland, 1991; Hart et al. 2013). The tagging programme on Sal similarly revealed that some turtles recorded as first-time nesters on Sal Island had previously been tagged elsewhere in Cabo Verde, indicating that non-resident individuals can enter the Sal nesting aggregation. We therefore quantified the seasonal proportion of non-resident individuals relative to the total number of tagged individuals to evaluate whether immigration from other Cabo Verdean nesting aggregations could explain a substantial proportion of the observed increase in nest counts.

### Tracking Loggerheads Nesting on Sal

To examine whether nest dynamics were associated with environmental changes at the foraging grounds, we tracked five adult females. Three Satellite-Relayed Data Loggers (SRDLs) manufactured by SMRU (Sea Mammal Research Unit, St. Andrews, UK) were deployed in 2011 (Roman-Torres et al. 2026), and two Splash 203 devices manufactured by Wildlife Computers were deployed in 2021. After oviposition, turtles were placed in a bottomless box to allow them to settle. Before device attachment, the carapace was cleaned of epibionts. Devices were then secured to the carapace using a two-part epoxy resin (Pure-2K, Power Fasteners, New Rochelle, NY, U.S.A., Schofield et al. 2010). Those five devices provided tracking data from October 2011 to February 2012 and from September 2021 to May 2022. We used these deployments, conducted ∼10 years apart, to assess whether the main post-nesting foraging areas remained consistent during the period of increasing nest abundance. A shift in foraging grounds could confound correlations between environmental conditions, demographic change and nest trends. Location data were managed using the Satellite Tracking and Analysis Tool (STAT, www.seaturtle.org), with filtering steps applied to exclude low-quality data (Roman-Torres et al. 2026). In brief, all GPS coordinates of Class B and 0 quality were removed, together with those obtained from fewer than 3 satellites. A speed filter was also applied to remove unrealistic displacement (>2 m s^-1^). This resulted in a dataset of 4,417 locations for five turtles.

### Collection of Environmental Parameters

To examine whether changes in oceanographic conditions could contribute to variation in nest counts, we extracted environmental variables from the identified foraging area. Two key ocean variables potentially influencing sea turtle reproduction, and therefore nest counts, were considered: chlorophyll-*a* concentration (Chl-*a*), a proxy of ocean productivity (Behrenfeld and Falkowski 1997) that has been linked to reproductive output in sea turtles (Ramírez et al. 2021; Valverde-Cantillo et al. 2019), and elevated sea surface temperature (SST), which can influence reproductive output through effects on physiological processes and nesting frequency (Nugraha et al. 2026; Weber et al. 2011).

Sea surface temperature and chlorophyll-a concentration were retrieved for both the foraging (11.12°–18.31° N, -14.50°–-20.66° W) and Sal breeding grounds (16.5°–16.9° N, -22.7°–- 23.1° W) from the OSTIA dataset (<u>Copernicus Marine Service</u>) and MODIS (<u>NOAA</u>) databases, respectively. Mean SST and cumulative Chl-a were calculated for 1–4 years before each nesting year to account for the variable remigration intervals of loggerheads (Cassill 2021; Nugraha et al. 2026; Phillips et al. 2014; Vander Zanden et al. 2015). For example, the 1-year mean SST for the 2008 season covered November 2007 to May 2008, because from June females migrate to Cabo Verde and remain there until nesting ends. For each nesting year, mean SST and cumulative Chl-a at the breeding grounds were calculated across the nesting months. Cumulative Chl-a was used to represent the integrated exposure to ocean productivity over the relevant period rather than productivity during a single month, better linking the accumulation of energy reserves in this capital breeder species.

Historical air temperature data for Sal Island were retrieved from NOAA’s Global Historical Climatology Network (GHCN) and Climate Anomaly Monitoring System (CAMS; <u>NOAA</u>), covering data from 1948 onwards. Because sea turtles nesting in a given year are likely to have hatched between 22 and 40 years earlier (Avens et al. 2015), we retrieved historical air temperatures using time lags reflecting this generation time.

### Aerial Survey and Breeding Sex Ratio

Under a temperature-driven sex-ratio bias hypothesis, we would expect a strongly female-biased breeding sex ratio (i.e. proportion of adult males and females that could contribute to the breeding pool) during the breeding season. Hence, to assess the breeding sex ratio, DJI Mavic 2 drones were flown over two nesting sites on Sal Island, Murdeira Bay and Costa Fragata beach (Fig. 1C) in two different nesting seasons, 2022 (April to October) and 2023 (May to September). These sites are known aggregation areas for adult turtles during the nesting season, with no records of juveniles or subadults. Drones were flown at an altitude of 60 m, providing a 100 m field of view at a speed of 10 m s^-1^, weather permitting (wind speed <11.2 m s^-1^; Dickson et al. 2022). Standardized transects were set parallel to the shore, at distances of 100–800 m offshore, ensuring continuous ocean coverage over a 5 km coastal stretch. Flights were conducted approximately every 13 days, thereby covering variation in breeding aggregation composition throughout the nesting season. We recorded videos, which also enabled the use of individual movement patterns to improve turtle detection.

In adults, males can be identified from the presence of a long tail (Schofield et al. 2017). The proportion of males and females detected through drone surveys was therefore used as an estimate of the breeding sex ratio, representing individuals present within breeding aggregations during the reproductive season. To minimize bias in detection and sex assignment, frame-by-frame analyses of drone footage incorporated strict image-quality controls. Each detected turtle was assigned a quality score from 1 (highest) to 5 (lowest), and only individuals scored 1–3 were included in the analysis. At these quality levels, the head, carapace, at least some flippers, and the posterior body/tail region were sufficiently visible to allow reliable sex assignment. Each detected turtle was assessed by two observers. Body measurements were also obtained for each identified turtle. Individuals falling outside the size range recorded for nesting adults (<63 cm straight carapace length, Nugraha et al. 2026) were classified as non-reproductive and excluded from the analysis, although no such individuals were detected on the videos. To assess whether the observed breeding sex ratio differed from parity (1:1), chi-square goodness-of-fit tests were performed separately for the 2022 and 2023 nesting seasons.

### Population Models for Sal Nesting Population

To investigate potential mechanistic links among historical air temperature, sex-ratio bias, and nest counts, we developed an eco-physiological individual-based model incorporating air temperature, hatchling sex ratio and survival rate. The aim of this model was to test whether warming-driven changes in sex ratios and recruitment could generate population trajectories consistent with the observed increase in nest counts. As our earliest nesting record dates to 2008, and females nesting in that year may have hatched approximately 22–40 years earlier, we used historical air temperature data from 1968, starting 40 years before the first nesting record. Air temperatures were transformed into sand temperatures using the following Sal-specific equation (Laloë et al. 2014) identified from buried temperature loggers at nest depth:

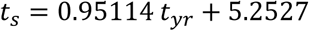

Where *t_s_* is the sand temperature (°C) and *t_yr_* is the mean air temperature from August-September of the given year (°C), representing the peak of the incubation season. The resulting temperature was then used to determine the sex ratio of each egg clutch using the following equation, taken from Laloë et al. (2014), following Girondot (1999):

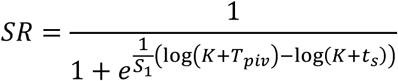

where SR is the proportion of males, *S*_1_ = −0.0336281 is the curvature parameter for the thermal performance curve, *K* = 0.1 is a constant, *T_piv_* = 28.95°C is the pivotal temperature and *t_s_* is the temperature experienced by that egg. We also incorporated metabolic heat produced within the clutch, with 25% of eggs in a nest experiencing a 0.5°C increase in temperature (Laloë et al. 2014).

Before applying the historical temperature trajectory, a 250-year burn-in period was included to allow the simulated population structure and sex ratio to reach equilibrium before applying the historical air temperature trajectory, ensuring the main simulation started from a realistic population, rather than from arbitrary initial conditions. During the burn-in period, the model used the mean temperature from the first decade of historical air temperature from Sal Island to establish initial sex ratios. In this case, the burn-in air temperature was 25.71°C (translating to a sand temperature of 29.71°C), which resulted in an initial population-wide sex ratio of 29% males. For comparison, we ran a null model in which temperatures varied around this baseline mean but did not include the observed long-term warming trend, allowing the contribution of warming-induced sex-ratio changes to population trajectories to be assessed.

A year in the model consists of three stages: reproduction, survival and development. Sexually mature males are assumed to appear every year, while adult females typically reproduce at intervals of 1–4 years, although longer intervals can occur, so only a fraction is present each year (Hays et al. 2010, 2014; Roman-Torres et al. 2026). The reproductive interval is randomly sampled from a normal distribution *N*(*μ* = 3, *σ* = 1) in discrete intervals after every reproductive event, as there is no evidence of individual interannual consistency.

At the start of the reproductive season, each female present (as determined by her reproductive interval) is assigned a number of nests, drawn from a normal distribution *N* (*μ* = 3, *σ* = 1) rounded to integers, based on clutch frequencies observed on Sal (Nugraha et al. 2026). Each nest, in turn, contains a number of eggs, randomly sampled from a Poisson distribution P (λ = 80, Nugraha et al. 2026). Demographic parameters used in the model (Supplementary Table S1) were based on a combination of published demographic information, biological assumptions, and model calibration. The parameters were calibrated so that the simulated population remained stable, producing nest counts comparable to those observed at the start of the time series in 2008. Because demographic data prior to 2008 were unavailable, this calibration assumes that the population was close to equilibrium at the beginning of the observed period. The juvenile stage included individuals <1 year old. From age 30, subadults matured with an annual probability of 35% so maturation within a cohort was spread over several years. Adult survival was set at 95%, consistent with the high survival expected for long-lived sea turtles and with values used in loggerhead demographic assessments (Conant et al. 2009). Note that the trend in overall population size was highly sensitive to the survival of subadults.

### Global Loggerhead Trends and Temperatures

Global loggerhead nesting trends were obtained from published and grey literature, covering 28 other populations (total populations = 29, including our data from Sal Island, Table 1) across nine Regional Management Units (RMUs), as defined by Wallace et al. (2023, Table 1). The coordinates and nesting months for each population are provided in Supplementary Table S2. For consistency, datasets using non-nest counts (e.g., female nester numbers) were standardized (Supplementary Table S3). Missing numerical values from graphical data were extracted using WebPlotDigitizer v4.6 (https://automeris.io/WebPlotDigitizer.html).

**Table 1.** Global loggerhead population data with sources and time series range. . Site coordinates and nesting months are provided in Supplementary Table S2. RMUs: Regional Management Units.

| Country/ Territory | Location | RMUs | Type of data | Time series period | Length of time series (years) | References |
|---|---|---|---|---|---|---|
| Australia | Gnaraloo Bay | Southeast Indian | Nest count | 2008 - 2015 | 8 | Thomson et al. 2016 |
| Australia | Woongarra | South Pacific | Individual nester count | 1988 - 2004 | 17 | Chaloupka et al. 2008 |
| Australia | Wreck Island | South Pacific | Individual nester count | 1988 - 2004 | 17 | Chaloupka et al. 2008 |
| Brazil | Bahia and Espírito Santo | Southwest Atlantic | Nest count | 1988 - 2003 | 16 | Marcovaldi and Chaloupka 2007 |
| Brazil | Rio de Janeiro, Farol de São Tomé | Southwest Atlantic | Nest count | 1992-2010 | 19 | Lima et al. 2012 |
| Cabo Verde | Sal Island | Northeast Atlantic | Nest count | 2008 - 2023 | 16 | This study |
| Cyprus | Alagadi | Mediterranean | Nest count | 1995 - 2019 | 25 | Omeyer et al. 2021 |
| Cyprus | Polis/Limni, Chrysochou | Mediterranean | Nest count | 1999 - 2015 | 17 | Casale et al. 2020 |
| Greece | Peloponnesus, Kyparissia | Mediterranean | Nest count | 1988 - 2024 | 37 | Margaritoulis et al. 2025 |
| Greece | Zakynthos Island, Laganas Bay | Mediterranean | Nest count | 1988 - 2021 | 34 | Margaritoulis et al. 2022 |
| Japan | Wakayama Prefecture, Minabe, Senri beach | North Pacific | Nest count | 1990 - 2001 | 12 | Hatase et al. 2002 |
| Japan | Shikoku island, Kamoda | North Pacific | Number of landings | 1988 - 2004 | 17 | Chaloupka et al. 2008 |
| Japan | Yakushima island, Maehama | North Pacific | Nest count | 1989 - 2015 | 27 | Martin et al. 2020 |
| Japan | Yakushima island, Inakahama | North Pacific | Nest count | 1988 - 2015 | 28 | Martin et al. 2020 |
| Japan | Yakushima island, Yotsusehama | North Pacific | Nest count | 1999 - 2015 | 17 | Martin et al. 2020 |
| Mexico | Quintana Roo state, Akumal | Northwest Atlantic | Nest count | 1995 - 2018 | 24 | González et al. 2020 |
| New Caledonia | La Roche Percée | South Pacific | Nest count | 2006 - 2019 | 14 | Barbier et al. 2023 |
| Oman | Masirah Island | Northwest Indian | Nest count | 2008 - 2016 | 9 | Willson et al. 2020 |
| South Africa | Maputaland Marine Reserve | Southwest Indian | Nest count | 1988 - 2010 | 23 | Nel et al. 2013 |
| Turkey | Antalya province, Kemer district, Çıralı | Mediterranean | Nest count and estimates | 1994 - 2020 | 27 | Casale and Margaritoulis, 2010 ; Sönmez et al. 2021 |
| United Kingdom Overseass Territory | Cayman Islands | Northwest Atlantic | Nest count | 1998 - 2020 | 23 | Cayman Islands Department of Environment 2022 |
| United States of America | Canaveral National Seashore | Northwest Atlantic | Nest count | 1988 - 2003 | 16 | Antworth et al. 2006 |
| United States of America | Dry Tortugas | Northwest Atlantic | Nest count estimates | 2001 - 2010 | 10 | Richards et al. 2011 |
| United States of America | Florida panhandle | Northwest Atlantic | Nest count | 1997 - 2022 | 26 | Florida Fish and Wildlife Conservation Commission |
| United States of America | Florida peninsula | Northwest Atlantic | Nest count | 1989 - 2022 | 34 | Florida Fish and Wildlife Conservation Commission |
| United States of America | Georgia | Northwest Atlantic | Nest count | 1989 - 2022 | 34 | Georgia Department of Natural Resources 2023 |
| United States of America | North and South Carolina | Northwest Atlantic | Nest count estimates | 2001 - 2010 | 10 | Richards et al. 2011 |
| - | Greater Caribbean | Northwest Atlantic | Nest count estimates | 2001 - 2010 | 10 | Richards et al. 2011 |
| - | Northern Gulf of Mexico | Northwest Atlantic | Nest count estimates | 2001 - 2010 | 10 | Richards et al. 2011 |

Historical air temperatures were retrieved using the same approach as for Sal Island, with temperatures extracted from 22 to 40 years before each nesting season to account for delayed recruitment into the breeding populations. These analyses were designed to test whether the relationship between historical temperature and subsequent nesting trends observed in Cabo Verde was detectable across loggerhead populations globally. Temperature trends for all sites are provided in the Supplementary Fig. S1 and nest count trends for all populations are shown in Supplementary Fig. S2.

### Statistical Analyses

Linear regression models tested the correlation between nest counts from Sal Island and (1) SST at the foraging ground, (2) Chl-a at the foraging ground, (3) SST and Chl-a interaction, (4) historical air temperature with time lags of 22 to 40 years, and (5) the interaction between historical air temperatures at the best-supported time lag and both SST and Chl-a. Model selection was performed using the Akaike Information Criterion (AICc) via the *AICcmodavg* package (Mazerolle 2023). No significant relationships were found for SST/Chl-a or their interactions, or the interaction between SST/Chl-a and temperatures from the best time lag. The strongest model correlated nest counts with historical air temperatures from the best time lag alone (Supplementary Table S4).

We then extended the correlation tests between historical air temperatures at time lags of 22 to 40 years and nest counts across 28 other loggerhead populations. These analyses tested whether relationships between historical temperatures and subsequent population trends observed in Cabo Verde are detectable across loggerhead populations globally, since there are no direct estimates of sex ratios in these populations. From these tests, we estimated the slopes of the relationships between population trends and historical air temperature, hereafter referred to as the Population Thermal Response (PTR). Where multiple significant time lags were identified, we used the PTR value from the correlation with the highest adjusted R². We also calculated the mean PTR (using the highest adjusted R²) and assessed its relationship with: (1) latitude, (2) temperature fluctuation and (3) long-term temperature trend.

Temperature fluctuation was calculated as the standard deviation of residuals from the linear regression of annual temperature against time, from 40 years before the first nest count to 22 years before the final nest-count record. Temperature trend was defined as the slope of the linear regression of annual temperature against time over the same period. Correction for multiple comparisons used the Benjamini-Yekutieli false discovery rate method (Narum, 2006). All statistical analyses were conducted using R version 4.2.2 (R Core Team 2022).

## Results

### Nest Count Trend of Loggerheads in Cabo Verde

In the early years of monitoring on Sal Island, only a few hundred nests were recorded annually but, around 2017, a rapid population increase was observed (Fig. 1D). This significant increase could theoretically stem from a large influx of new migrants from other nesting populations. However, using a mark–recapture approach with PIT tagging, we could identify turtles nesting on Sal Island that were tagged elsewhere in Cabo Verde, revealing non-Sal-resident individuals. The average proportion of non-resident turtles was 4.57%, ranging annually from 1.42% (in 2009) to 8.86% (in 2021). This suggests that, although there is some influx of individuals from other Cabo Verdean aggregations, the proportion is very low and therefore cannot drive the dramatic increase in nest counts observed.

### Oceanic Conditions and Nesting Dynamics

Using Chl-a retrieved from <u>NOAA</u>, we found no significant correlation between nest counts and Chl-a levels at either the foraging ground (Fig. 2A) or the breeding site (Fig. 2C), suggesting that food availability is not a primary driver of the observed increase in nest counts (Fig. 2B, LM: foraging ground adjusted R² = -0.06, *p* = 0.759; Fig. 2D, breeding ground adjusted R² = -0.06, *p* = 0.74). No significant correlations were detected either when investigating different timelines for remigration intervals (results for 1 to 4 years are shown in Supplementary Fig. S3).

**Fig. 2:**
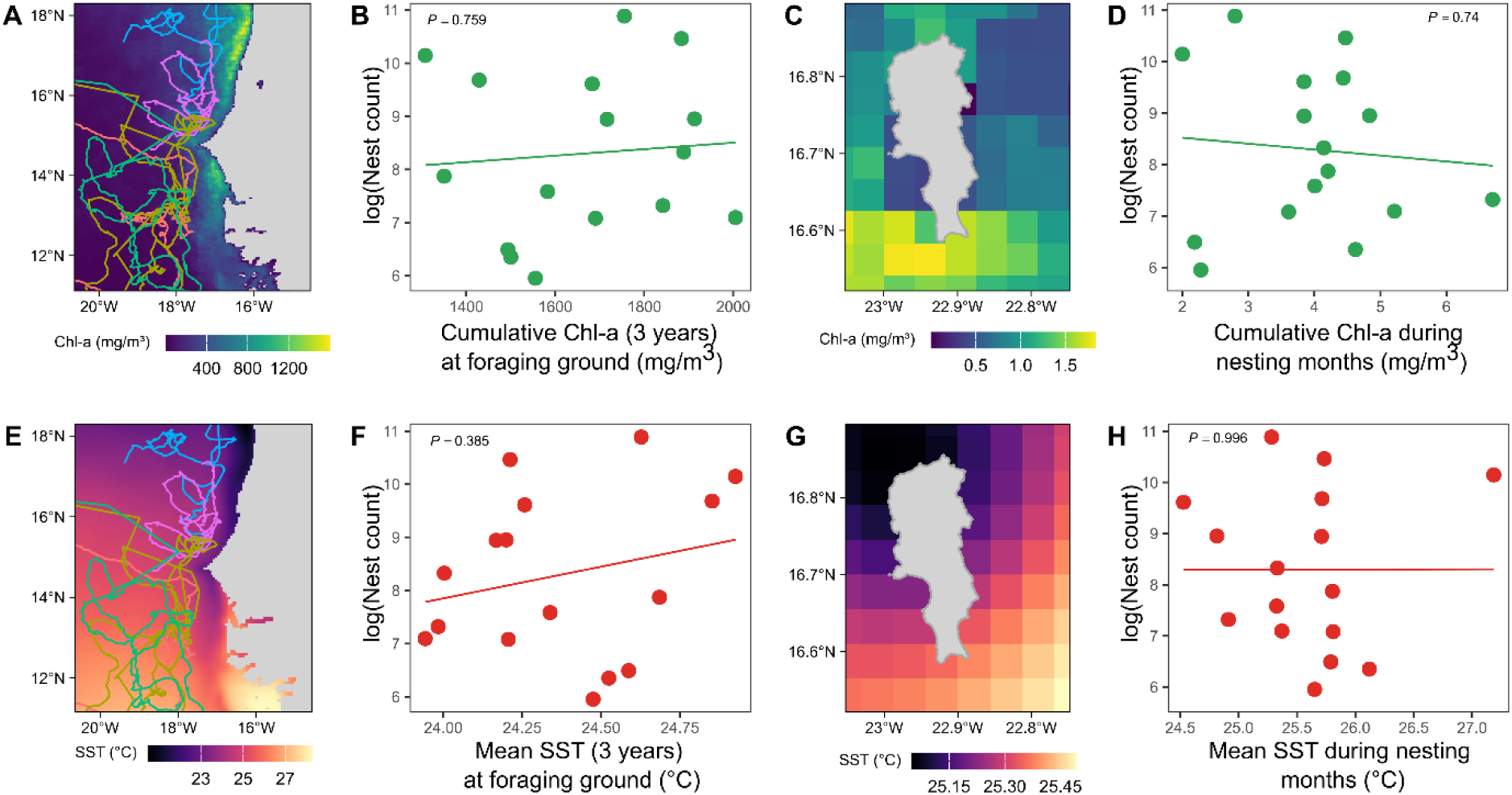
Relationship between ocean environmental parameters and nest counts of loggerhead turtles on Sal Island, Cabo Verde. **A.** Cumulative chlorophyll-a concentration (Chl-a, mg m^-3^) in the foraging grounds. Coloured lines indicate the post-nesting trajectories of five adult females tracked via Argos satellite telemetry in 2011 (n = 3) and 2021 (n = 2). **B.** Relationship between 3-year cumulative Chl-a in the foraging ground and nest counts. **C.** Cumulative Chl-a in the breeding ground during the breeding season. **D.** Relationship between cumulative Chl-a in the breeding grounds and nest counts. **E.** Mean sea surface temperature (SST, °C) and turtle trajectories in the foraging grounds. **F.** Relationship between 3-year mean SST in the foraging ground and nest counts. **G.** SST in the breeding ground during the breeding season. **H.** Relationship between SST in the breeding ground and nest counts.

SST at the breeding ground (Fig. 2G) showed no significant relationship with nest counts (Fig. 2H, LM, adjusted R² = -0.07, *p* = 0.996). Similarly, no correlations were detected between SST in the foraging ground (Fig. 2E) and nest counts (Fig. 2F, LM, adjusted R² = -0.01, *p* = 0.385, different lag years are provided in Supplementary Fig. S3).

### Historical Air Temperatures and Sex Ratios

Our main hypothesis is that the observed increase in nest counts is explained by shifts in primary sex ratio towards females driven by rising air temperatures, and, by extension, incubation temperatures. Between 1968 and 2023, spanning approximately two sea turtle generations, air temperatures during the core nesting period increased by more than 2.5 °C in Cabo Verde (Fig. 3A, linear model (LM), adjusted R² = 0.81, *p* < 0.001).

**Fig. 3:**
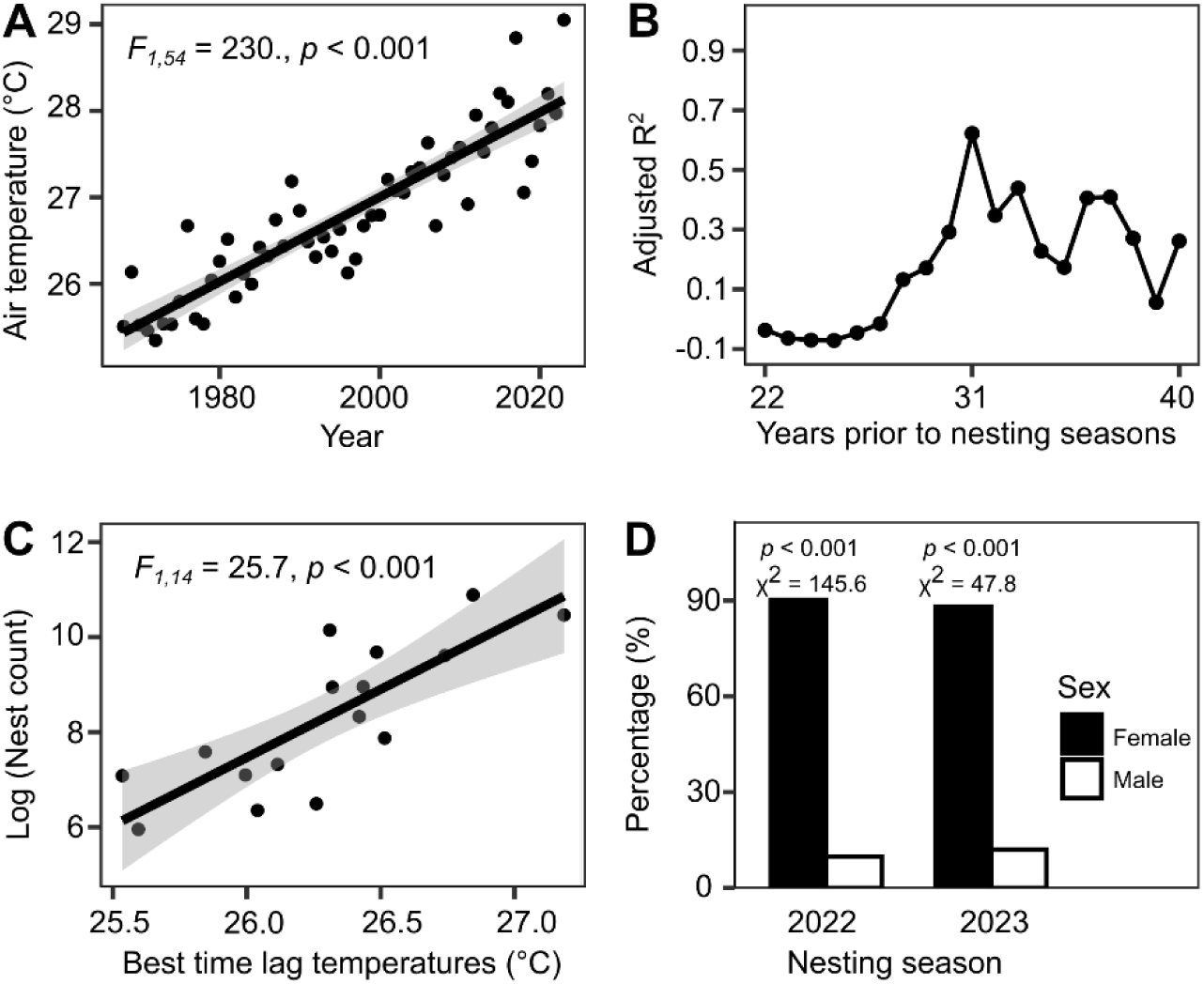
Correlation of loggerhead nest counts on Sal Island with historical air temperatures and detected breeding sex ratio. **A.** Air temperature trend during the core incubation period (August-September for the 1968-2023 time period). **B.** Adjusted R^2^ values from linear regressions of log-transformed nest counts against historical air temperatures, assuming time lags of 22 to 40 years (reflecting estimated age at maturity). The strongest correlation occurs with a 31-year time lag. **C.** Relationship between historical air temperatures 31 years prior to the nesting seasons and nest counts. **D.** Breeding sex ratio inferred from drone monitoring were significantly female-biased in both the 2022 (203 females and 22 males) and 2023 (73 females and 10 males) nesting seasons.

We evaluated the correlation between annual nest counts from 2008 to 2023 and air temperature at time lags of 22–40 years, reflecting the likely birth years of nesting females. Positive correlations were observed for lags of 28 to 40 years between nest count and historical air temperatures (Fig. 3B), with the strongest correlation corresponding to a 31-year time lag (Fig. 3C, LM, adjusted R^2^ = 0.62, *p* < 0.001).

### Current Breeding Sex Ratio

A total of 308 turtles from drone footage were sexed based on the presence/absence of an elongated tail. Overall, 90% of individuals were classified as females (Fig. 3D), with 203 females and 22 males observed in 2022 (90.2% females) and 73 females and 10 males identified in 2023 (88.0% females). Breeding sex ratios were significantly female-biased in both years (2022: χ²(1) = 145.6, *p* < 0.001; 2023: χ²(1) = 47.8, *p* < 0.001). Monthly female-to-male ratios varied from 0 to 17 females per male in 2022 and from 0 to 16 in 2023, but were female-biased in all months when both sexes were observed. Despite this monthly variation, there was no obvious seasonal shift in the breeding sex ratio on Sal Island, as male and female abundance broadly peaked in parallel, with few males appearing to remain resident throughout much of the season (Supplementary Fig. S4).

### Population Dynamics Model for Sal Nesting Population

To investigate potential mechanistic links among historical air temperature, sex-ratio bias and nest counts, we developed a population dynamics model incorporating sand temperature, hatchling sex ratio and survival rate (Fig. 4A). When using historical sand temperature data, the model predicted increasing trends in nest counts and a positive correlation between nest counts and air temperature (Fig. 4B, C), along with a progressive decline in the breeding sex ratio with fewer males and more females in each life stage (Fig. 4D). When generational time lags were considered, these simulated trends corresponded with the real, observed pattern of temperature-related population increase seen in Sal. Similarly to our observed data, adjusted R² values were highest for time lags ranging from 30 to 35 years (Supplementary Fig. S5A), with the highest correlation detected for a 32-year time lag. The null model that randomized sand temperatures did not reproduce this increase in nest counts observed in the very recent past (Fig. 4B) or primary and adult sex ratio shifts (Supplementary Fig. S5B).

**Fig. 4:**
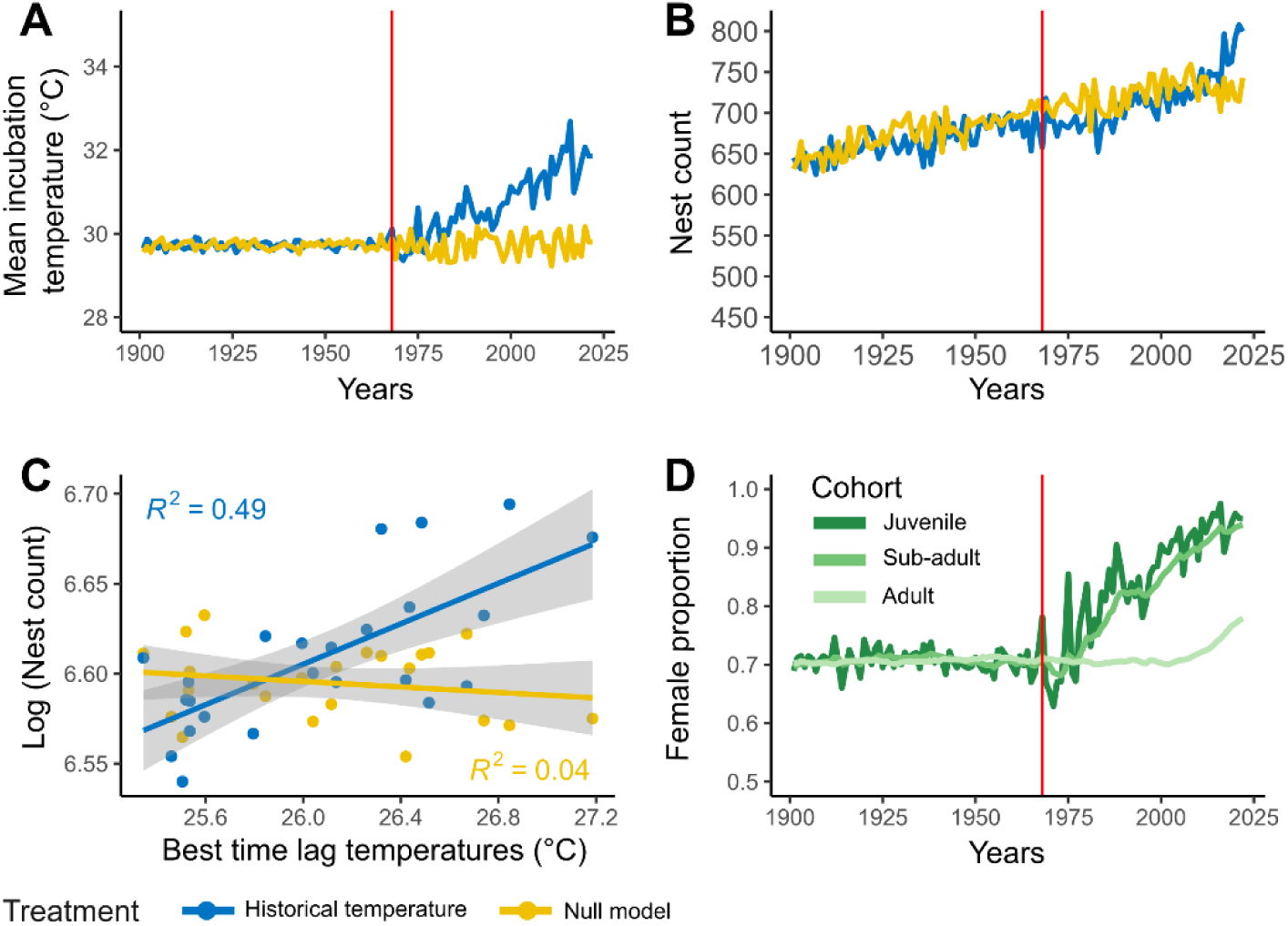
Simulated population response to increasing historical air temperatures on Sal Island. **A.** Sand temperatures from 1968 to 2022 were used as inputs for population dynamics simulations (blue) and the null model where observed temperatures were randomised (yellow). **B.** Predicted number of nests. **C.** Linear relationships between log-transformed modelled nest counts and temperature 32 years prior to the nesting season for both the historical air temperature-based and null models. **D.** Predicted cohort-specific sex ratios (female-based) in response to historical air temperatures on Sal Island. Red vertical lines indicate the end of the 250-year model burn-in period.

As expected, sex ratios in juvenile and subadult cohorts rapidly changed with incubation temperature, while adult cohort sex ratios reflect generational time lags and correlate with historical air temperatures. We further found the adult cohort to be female biased with a rate of about 80% by the end of the simulation, comparable to the observed ratio in our population (Fig. 4D). Overall, the congruence between our simulations and our empirical data for Sal Island suggests that the rapid population expansion of this TSD species is strongly associated with climate warming.

### Global Loggerhead Population and Historical Air Temperatures

If warming-driven feminisation shapes loggerhead demography generally, similar relationships should appear elsewhere, especially where warming has been strong. We therefore repeated the analyses linking nest counts to historical air temperatures for another 28 loggerhead populations globally. Among the 29 populations examined (including Sal Island), 13 displayed a significant positive relationship with historical air temperatures, with best-predicting time lags ranging from 22 to 40 years. The strongest positive relationships were detected across different geographic regions, including Cabo Verde, Mexico, and Cayman Islands. By contrast, three populations showed significant negative relationships with historical air temperatures namely the two Cypriot populations (Alagadi and Chrysochou) and Kyparissia, Greece (Fig. 5A, B; Supplementary Fig. S6). PTR (the slope of log nest count on lagged temperature) was context-dependent, showing a negative relationship with absolute latitude (Fig. 5C, LM, best PTR, adjusted R² = 0.31, F_1, 27_ = 13.9, *p* < 0.001), indicating that relationships between historical air temperature and nesting trends were more positive at lower latitudes and weaker or more negative at higher latitudes. We observed similar results when averaging the PTR across multiple significant time lags (Supplementary Fig. S7A, LM, mean PTR, adjusted R^2^ = 0.31, F_1, 27_ = 13.7, *p* < 0.001).

**Fig. 5:**
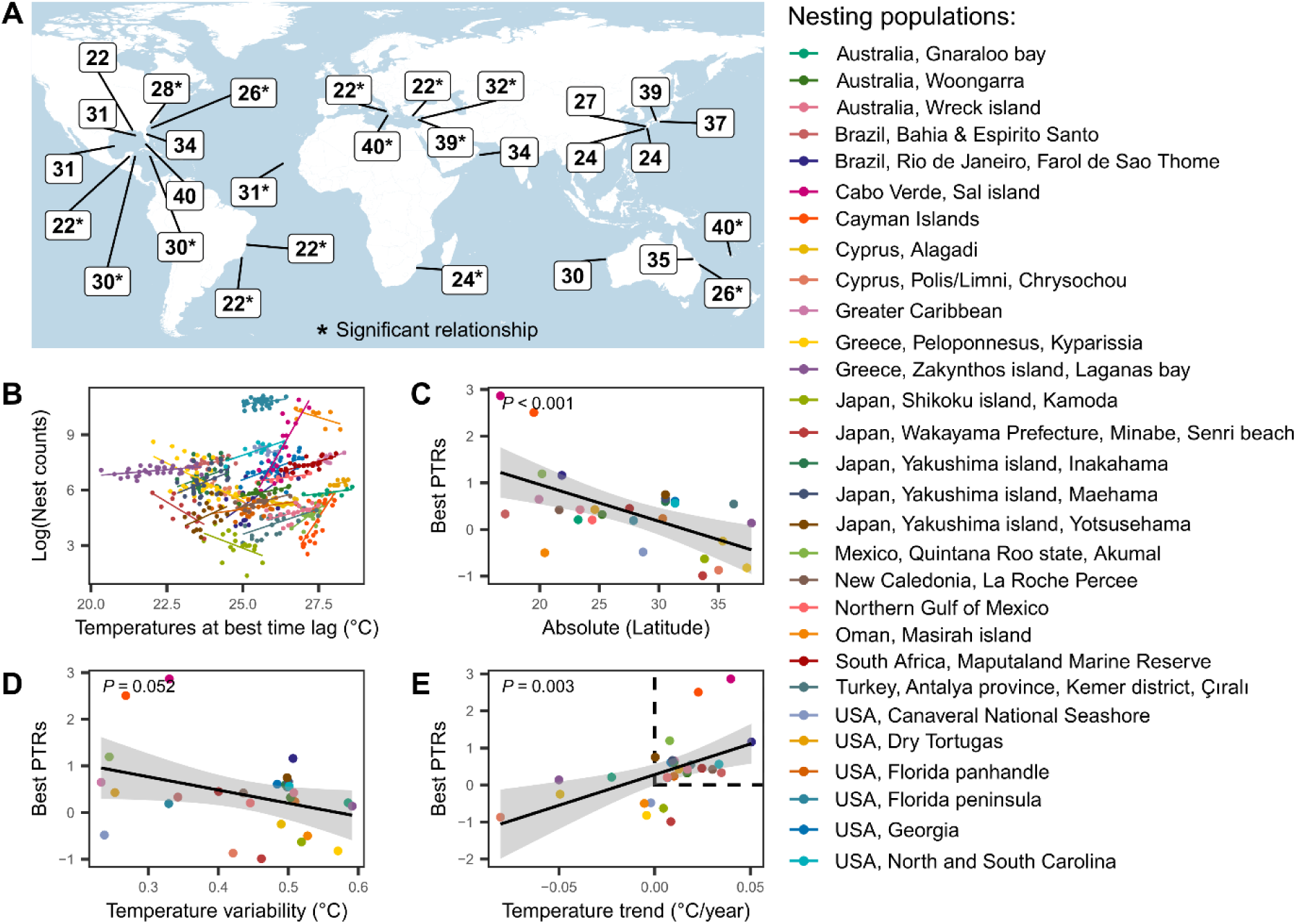
Global loggerhead population trends in relation to historical air temperatures. **A.** Best-predicting time lags (in years) for the relationship between loggerhead nest counts and historical air temperatures. Numbers indicate the time lag associated with the strongest temperature-nest count correlation. Numbers with an asterisk indicate a significant correlation. **B.** Regression lines illustrating the relationship between nest counts and historical air temperatures at the best-predicting time lag for each population. Colours represent different nesting populations. The predicted slope values from these relationships are termed the Population Thermal Response (PTR). **C.** Relationship between the strongest PTR and the absolute latitude of nesting sites. **D.** Relationship between strongest PTR and seasonal temperature variability (residuals around the annual temperature trend) during the nesting season. **E.** Relationship between the strongest PTR and the slope of the temperature trend at each site. Dashed lines show populations with positive PTR and exposed to warming trends.

We postulated that this geographic pattern may stem from temperature fluctuations or, alternatively, from the degree of warming across latitudes. We found that where temperature fluctuations were higher, population trends were less likely to correlate with historical air temperatures (Fig. 5D, LM, best PTR, adjusted R^2^ = 0.10, F_1, 27_ = 4.31 *p* = 0.052 after FDR correction; Supplementary Fig. S7B, LM, mean PTR adjusted R^2^ = 0.10, F_1, 27_ = 4.2, *p* = 0.05). Conversely, we found that PTR increased with the rate of historical warming (Fig. 5E, LM, best PTR, adjusted R^2^ = 0.26, F_1, 27_ = 11.9, *p* = 0.003; Supplementary Fig. S7C, LM, mean PTR, adjusted R^2^ = 0.25, F_1, 27_ = 10.1, *p* = 0.004), indicating that populations experiencing stronger warming tended to show more positive relationships between historical air temperature and subsequent nesting trends.

## Discussion

### Warming, sex-ratio bias and demographic imbalance

Global warming is impacting species in unprecedented ways (Bellard et al. 2012; Poloczanska et al. 2013). For taxa with temperature-dependent sex determination (TSD), these impacts are not only physiological but also demographic (Mitchell et al. 2010), because incubation conditions shape the composition of future breeding populations. Here, we provide multiple lines of evidence that warming has contributed to a strongly female-biased breeding population and rapid nest count growth in one of the world’s largest loggerhead turtle populations. The temperature-demography relationship further explains population trends globally for loggerhead turtles. The apparent conservation success of population expansion may partly reflect the maturation of female-biased cohorts and an emerging form of demographically imbalanced recovery, rather than a healthy population rebound. This pattern illustrates how climate warming can promote population growth in some regions of the world, while potentially creating longer-term risks to reproductive resilience (Mazaris et al. 2017; Patrício et al. 2021).

On Sal Island, nest counts have risen roughly 100-fold since 2008, with the sharpest growth occurring after 2017 and reaching over 50,000 nests in 2021. This surge is primarily driven by neophyte nesters (Hays et al. 2022). We found the best predictor of this nesting trend to be historical air temperature, with a generational lag of approximately 30 years, matching previous age-at-maturity estimates for loggerheads. Petitet et al. (2012), for example, estimated a mean age at maturation of 31.8 years, while Bjorndal et al. (2000) estimated 26.5 years to reach the minimum mature size and suggested that sexual maturity likely occurs closer to 30 years or older. Van Houtan and Halley (2011) adopted a 31-year lag based on estimated ages at first breeding of 30–32 years. Because survival is generally assumed to be similar between sexes (Hays et al. 2017), a parsimonious explanation is that climate-driven sex-ratio bias has contributed to increased recruitment and, consequently, increased nesting activity with more adult females (Hays et al. 2023; Laloë et al. 2024).

To determine the breeding sex ratio, we used drone surveys. They revealed a strongly female-biased breeding aggregation, with approximately 90% of individuals classified as females. This level of female bias is consistent with model-based studies of hatchling sex ratios in Cabo Verde, which predict female-biased production, including approximately 84% female hatchlings across the country (Tanner et al. 2019), 79% on Boa Vista Island (Abella-Perez et al. 2016), 77% on Santa Luzia Island (Rocha et al. 2015), and 70% on light-sand and 93% on dark-sand beaches on Sal Island (average: 81.5%, Laloë et al. 2014). Moreover, this level is also comparable with the extreme female-biased sex ratios reported in other sea turtle populations, such as the northern Great Barrier Reef green turtle population where females accounted for 99% of juveniles and 87% of adults (Jensen et al. 2018).

Several factors could influence the observed breeding sex ratio, including sex-specific survival, breeding periodicity, movement behaviour and detectability. However, none of these factors is likely to fully explain the magnitude of female bias observed here. Sex-specific mortality remains poorly quantified in sea turtles (Schofield et al. 2020), and equal survival between sexes is commonly assumed in demographic models (Chaloupka and Limpus 2005). In addition, males typically breed more frequently than females (Hays et al. 2014), which should, if anything, make the breeding aggregation less female-biased than the underlying adult population. Finally, surveys were conducted over breeding aggregations where males and females spatially overlap, and strict image-quality filtering reduced the likelihood of systematic bias in sex identification. While we acknowledge that no observational method is entirely free from potential detectability biases, such biases would need to be both strong and consistently directional to produce the level of female bias observed in our data (approximately 90% females). We also repeated the drone survey over two consecutive years to assess the consistency of the observed breeding sex ratios. Notably, the sex ratio estimated from drone surveys is based on the presence of mature males and females in the breeding aggregation. The breeding sex ratio would therefore be expected to be less female-biased than the adult sex ratio because reproductive periodicity differ between sexes: mature females generally breed at multi-year intervals, whereas males can return to breeding grounds more frequently (Hays et al. 2014). Taken together, these observations support the interpretation that recent recruitment has been influenced by warmer incubation conditions, resulting in a strongly female-biased breeding population.

### Limited evidence for alternative environmental drivers

To evaluate whether contemporary environmental conditions could explain the observed population trajectory in Sal, we examined relationships between nesting trends and oceanographic conditions at the foraging and breeding grounds. Neither sea-surface temperature (SST) nor ocean productivity (chlorophyll-a concentration) showed any significant relationship with nest counts, even when accounting for multi-year lags linked to female remigration intervals. This contrasts with findings from other regions (Chaloupka et al. 2008; Saba et al. 2007; Valverde-Cantillo et al. 2019). Such regional differences may reflect the oceanographic context of West Africa, where upwelling sustains consistently high productivity, punctuated by periodic cooling events (Benazzouz et al. 2014; Jamal et al. 2025). Thus, while fluctuations in food availability or SST within seasons may influence short-term remigration intervals, these factors are unlikely to explain the sustained increase in nest counts observed on Sal Island. It is also worth noting that this population experiences high bycatch levels, which may obscure correlations between population expansion and contemporary environmental variables (Cardona et al. 2025). Overall, historical air temperature remains the environmental variable most strongly associated with nesting trends in this population.

### Mechanistic support from population modelling

To provide additional evidence for a mechanistic link between warming, sex-ratio bias, and nesting trends, we developed an eco-physiological population dynamics model. Simulations using historical air temperature trajectories reproduced the increase in nest counts, whereas null models without directional warming did not generate comparable dynamics. Importantly, the model also projected a progressive decline in male representation within the adult population, reflecting cumulative effects of temperature-driven sex ratio shifts across cohorts.

As expected, sex ratios in juvenile and subadult cohorts responded rapidly to temperature variation, whereas adult sex ratios reflected generational time lags, with shifts becoming detectable after approximately 30–35 years (Avens et al. 2015; Bjorndal et al. 2000; Van Houtan and Halley 2011; Petitet et al. 2012). Under the model parameterisation, historical warming generated an approximately 20% increase in simulated nest counts relative to the no-warming scenario (which also shows a general upward trend). This demonstrates that historical warming alone can generate a measurable increase. For our model, we made assumptions around key parameters, including the pivotal temperature, the temperature that produces a 1:1 sex ratio within a clutch. Although we used a pivotal temperature previously applied to the Sal sea turtle aggregation (Laloë et al. 2014), it is possible that this value might not apply to our study population (Lockley and Eizaguirre 2021; Yen et al. 2026). Such variation may partly explain differences between model expectations and empirical counts and therefore warming could be associated with a larger contribution to population increase.

### Role of conservation efforts

Since our model does not predict the absolute magnitude of the increase in nest counts, it suggests conservation measures and other demographic factors also contribute to the upward trend. Cabo Verde is globally recognised as a stronghold for loggerhead turtles (Hays et al. 2022; Nugraha et al. 2026; Patino-Martinez et al. 2022; Roch et al. 2026; Taxonera et al. 2022), and ongoing conservation initiatives have likely reinforced population growth. Protection of nesting females through night patrols has lowered adult mortality on land. At sea, however, bycatch mortality remains substantial: industrial longline fisheries off northwestern Africa are estimated to kill thousands of loggerheads annually, including adults from the Cabo Verde population (Cardona et al. 2025; De La Hoz Schilling et al. 2023). Because comparable historical estimates of bycatch mortality are unavailable, we cannot determine whether this pressure has changed through time. This matters because simulated population trends were highly sensitive to subadult survival, the life stage most exposed to longline bycatch, so changes in at-sea mortality could have contributed to the observed increase in either direction. Conservation has nevertheless likely contributed to population recovery through reductions in anthropogenic mortality on nesting beaches.

However, conservation alone is unlikely to explain the totality of the demographic pattern observed here. First, the timing of the rapid increase in nest counts is more consistent with the expected maturation of cohorts produced during earlier warming than with the chronology of local conservation interventions. Second, if the recent influx of new recruits were primarily driven by hatchery-based production, a more balanced breeding sex ratio would probably be expected, particularly if hatcheries increased recruitment without strongly biasing incubation temperatures. Finally, if conservation were the sole driver of population increase, we would not expect population trends across global loggerhead populations, which have experienced different conservation histories, to show a geographic relationship with historical air temperatures. Our results therefore suggest that conservation efforts acted as a complementary driver of recovery, while warming-induced sex-ratio bias contributed substantially to the observed demographic signal altering the population’s adaptive potential (Eizaguirre and Baltazar-Soares 2014).

### Beyond local dynamics: a global demographic pattern

Extending our framework globally, we found that historical air temperature trajectories were associated with nesting trends across 29 loggerhead populations. The strongest relationships occur at low-latitude rookeries, where high baseline temperatures and limited seasonal variability may increase the persistence of warming effects on cohort sex ratios (Pike 2014; Witt et al. 2010). In these regions, sustained warming may therefore promote increasingly female-biased cohorts that coincide with population growth, while potentially contributing to demographic imbalance over longer timescales. By contrast, higher-latitude populations, where interannual temperature fluctuations are greater, show weaker correlations, suggesting that environmental variability may buffer against consistently female-biased cohort production, as observed in other TSD species such as snapping turtles (Leivesley et al. 2022). Importantly, sex ratios were not measured across all these populations. Rather, these relationships identify thermal regimes in which the demographic consequences expected from persistent warming may be more pronounced.

These results suggest that the combination of relatively stable temperatures and stronger warming trends at lower latitudes is more likely to generate conditions under which temperature-driven female recruitment could contribute to population increases. At higher latitudes, greater temperature variability may maintain a broader range of incubation conditions and thereby reduce persistent directional sex-ratio bias. Populations showing both a strong dependence of demographic trends on historical air temperatures and greater exposure to rapid warming may therefore be particularly vulnerable to future demographic imbalance (Fig. 5E, points within the dashed square).

To quantify the variation among populations in their demographic response to historical temperature, we developed the Population Thermal Response (PTR), a quantitative index linking long-term demographic trends to thermal history. The PTR captures variation among populations in the strength and direction of the relationship between demographic trends and historical temperature. It may serve as a useful screening framework to target aggregations where direct monitoring of sex ratios may be particularly informative for conservation. Populations with high positive PTR values, in which positive nesting trends are most positively associated with historical warming, may paradoxically face greater demographic risk due to sex ratio imbalance if persistent male scarcity begins to constrain mating and fertilisation. This coexistence of recovery and emerging demographic risk challenges conventional interpretations of conservation success and highlights the need to integrate physiological thresholds and sex-ratio dynamics into demographic projections (Fuentes et al. 2024; Hays et al. 2017).

### Implications for conservation and monitoring

An important implication of this study is that rapid increases in nesting abundance may signal emerging demographic imbalance in species with temperature-dependent sex determination. For sea turtles, while highly female-biased populations may initially exhibit strong population growth, long-term viability depends on sufficient male availability (Hays et al. 2017). If male scarcity becomes limiting, current recovery trajectories could be associated with reduced reproductive resilience.

Our findings highlight that population monitoring focused solely on abundance may be insufficient to assess population health in TSD species. Population growth can coexist with underlying demographic imbalance from sex ratios, and conservation assessments should therefore consider both abundance and demographic structure. More broadly, this study suggests that recovery in some TSD species under climate warming may involve complex demographic shifts, where an increase in abundance does not necessarily reflect long-term stability. This principle may extend beyond sea turtles to other TSD species, where climate-driven changes in offspring sex ratios could alter population trajectories before demographic constraints associated with sex-ratio imbalance become apparent.

## Data Accessibility

All data and R scripts used in this study are available at Zenodo repository https://doi.org/10.5281/zenodo.24088945

## Figure Sources and Image Credits

For figure 1A, the follwoings are the source and the credit of figures:

1. The person is the first author of the manuscript. Photo was taken by James O. Bazely.
2. The picture of the loggerhead turtle with a satellite tracking attached was taken by Samuel J. Shrimpton.
3. Satellite imagery was obtained from https://www.pexels.com/photo/gray-and-white-satellite-41006/. It was edited to remove any label/brand names on it.
4. Drone image was taken from https://www.dji.com/uk/support/product/mavic-2.
5. Loggerhead turtles on the map in is edited to remove any text on it. Source: https://www.fisheries.noaa.gov/species/loggerhead-turtle.

For figure 1D, the turtle image was created by ChatGPT on May 11, 2025 with the following prompt: “a loggerhead illustration facing to the right handside”.

## DECLARATIONS

## Funding

This research was funded by UK Research and Innovation Natural Environment Research Council NE/V001469/1 to C.E and J.T as well as NE/X012077/1 to C.E. Further funding came from Queen Mary University of London to C.E. Fellowship for F.A.D.N. (FR202403000977) is provided by Beasiswa Pendidikan Indonesia (Indonesian Education Scholarship, Center for Higher Education Funding and Assessment and Indonesian Endowment Fund for Education). Associação Projeto Biodiversidade was funded by U.S. Fish and Wildlife Service (USFWS), The Regional Partnership for Coastal and Marine Conservation in West Africa (PRCM).

### Acknowledgement

We are grateful to SOS Tartarugas, Associação Projeto Biodiversidade, and the volunteers for their continuous field monitoring of sea turtles on Sal Island. The authors also would like to thank the editor and two anonymous reviewers for their recommendations that improved the manuscript.

## Permits

Permits were granted by the Cabo Verde Government through the partnership agreement between Associação Projeto Biodiversidade and the Ministry of Agriculture and Environment. For this study, members of the Eizaguirre lab operated under permit numbers 215/DNA/2024 and 474/DNA/2025.

## Author contributions

C.E designed the study and supervised the overall project. K.F, A.L, and A.T organised field patrols. S.M.C, A.T, C.E organised field work. All coauthors contributed to field work. B.R., A.T. and C.E organised and conducted drone surveys. L.S. analysed drone data. F.A.D.N supervised and conducted data analyses with support from C.E and J.G. R.A and J.T performed the simulation. F.A.D.N wrote the manuscript with contributions from C.E and S.J.R. All authors contributed to and approved the final version of the manuscript.

