## Supplementary figure S1 to S7 for "Warming-induced sex bias underlies rapid recovery but may generate demographic imbalance in sea turtles"

**Running title: Climate warming and sea turtle recovery**

Fitra Arya Dwi Nugraha[1], Rebekka Allgayer[2], James Gilbert[1], Laura Sivess[1], Kirsten Fairweather[3], Artur Lopes[3], Sandra M. Correia[4], Berta Renom[3], Albert Taxonera[3], Stephen J. Rossiter[1], Justin Travis[5,6], Christophe Eizaguirre[1]

[1] Department of Biology, School of Biological and Behavioural Sciences, Queen Mary University of London, London E1 4NS, United Kingdom

[2] RSPB Centre for Conservation Science, The Lodge, Sandy SG19 2DL, United Kingdom

[3] Associação Projeto Biodiversidade, Santa Maria 4111, Sal, Cabo Verde

[4] Instituto do Mar, Cova de Inglesa, CP 132, Mindelo, Cabo Verde

[5] Department of Biosciences, Durham University, DH1 3LE Durham, United Kingdom.

[6] Faculty of Forestry, Universitas Gadjah Mada, Jl. Agro No. 1 Bulaksumur, Sleman, Yogyakarta, Indonesia

<u>**Supplementary Figure S1:**</u>

**Air temperature trends during the nesting months across global loggerhead nesting sites.** Trends were estimated using linear regression, with mean air temperature during the nesting months shown on the y-axis and year on the x-axis. For each of the 29 populations, the first point represents air temperature 40 years before the first recorded nest count in the time series, while the final point represents air temperature in the year of the most recent recorded nest count.

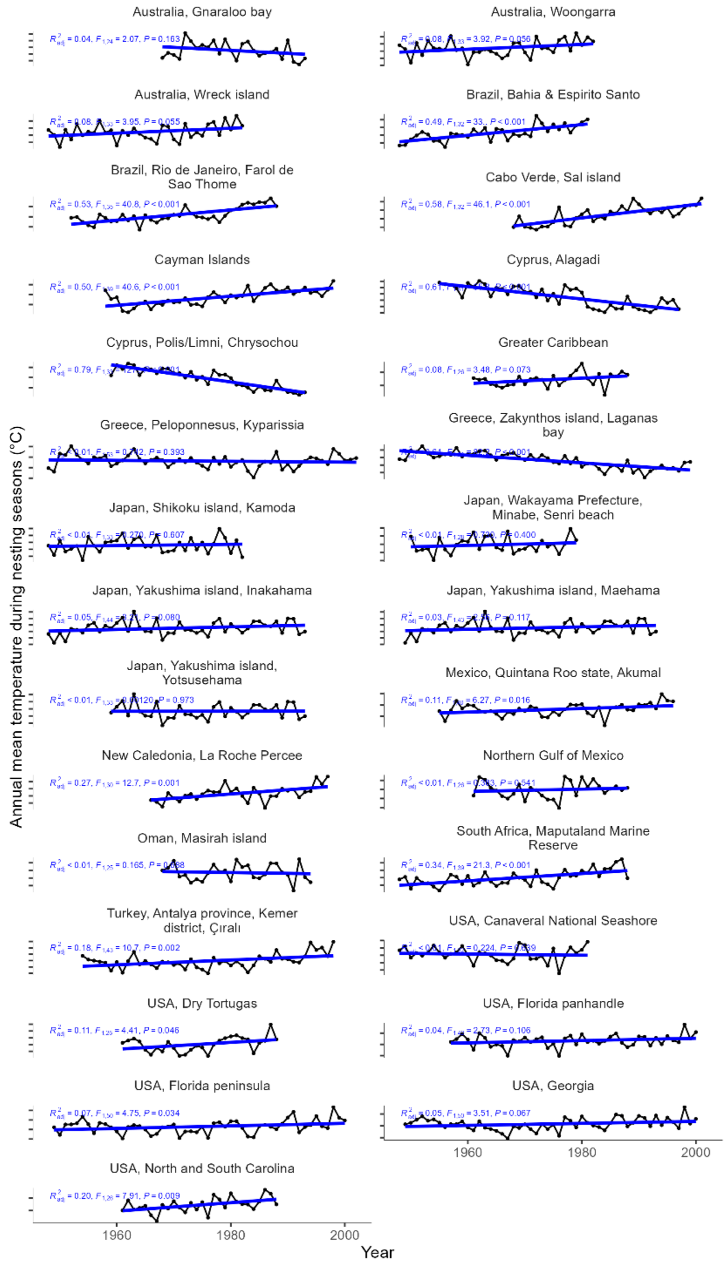

**Supplementary Figure S2:**

**Global loggerhead population trends estimated using linear regression models.** The y-axis represents log-transformed nest counts, and the x-axis represents year. Three populations showed declining trends: (1) Kamoda, Japan; (2) Senri Beach, Japan; and (3) Masirah Island, Oman. USA = United States of America.

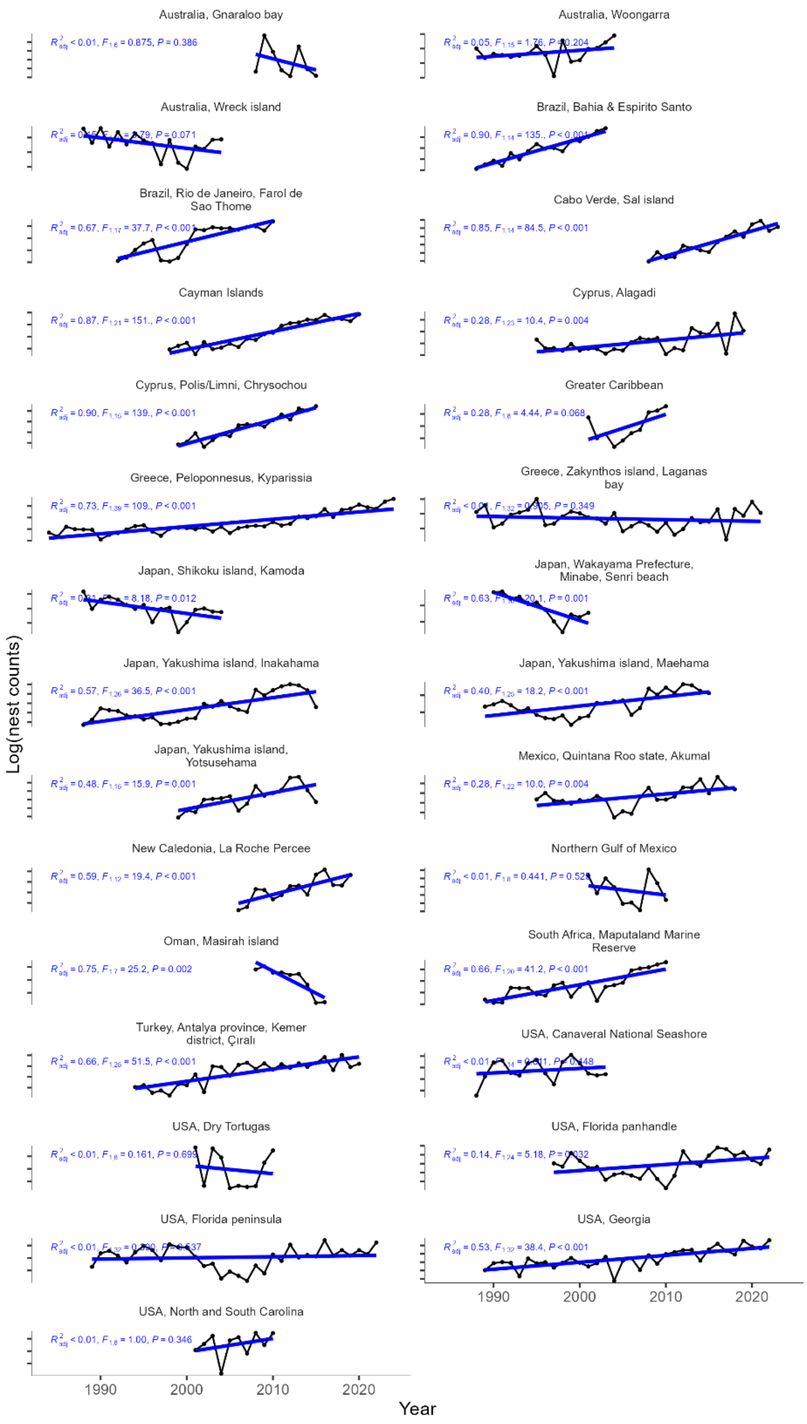

**<u>Supplementary Figure S3:</u>**

**Linear relationships between log-transformed nest counts and two oceanographic variables, sea surface temperature (SST) and chlorophyll-a concentration (Chl-a), at both breeding and foraging grounds for loggerhead turtles nesting in Sal Island.** At foraging grounds (fg), mean SST and cumulative Chl-a were calculated using time lags of one to four years. At breeding grounds (bg), mean SST and cumulative Chl-a were calculated for the nesting season (June–October).

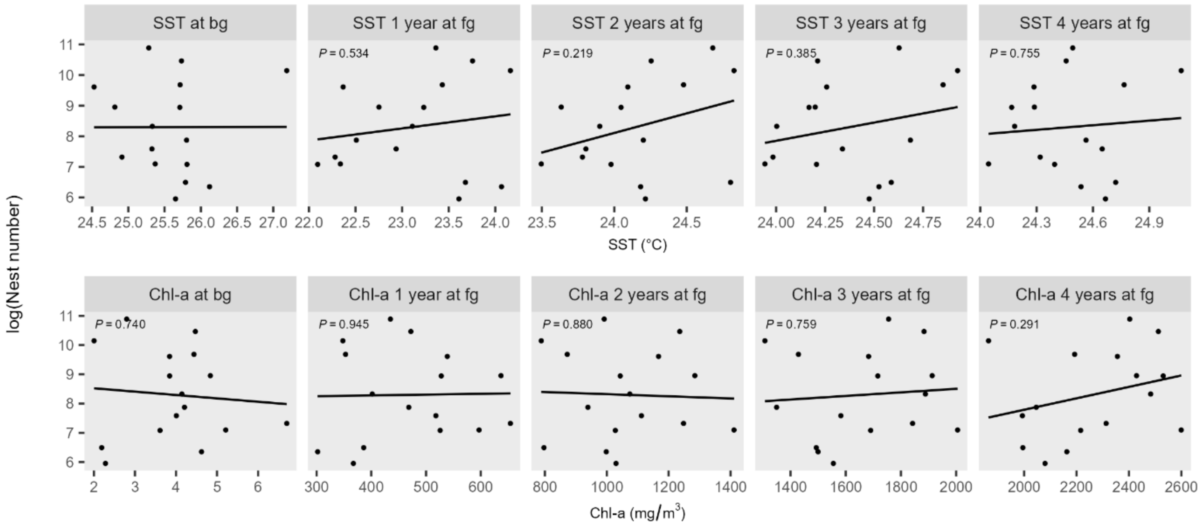

**Supplementary Figure S4:**

**The dynamics of breeding sex ratio surveyed using drones during the 2022 and 2023 nesting seasons at the breeding ground.** (A and B) Average monthly number of sexed individual turtles per survey, calculated by dividing the number of individuals recorded by the number of surveys conducted in each month. (C and D) Monthly contribution (%) of each sex relative to is total annual detections.

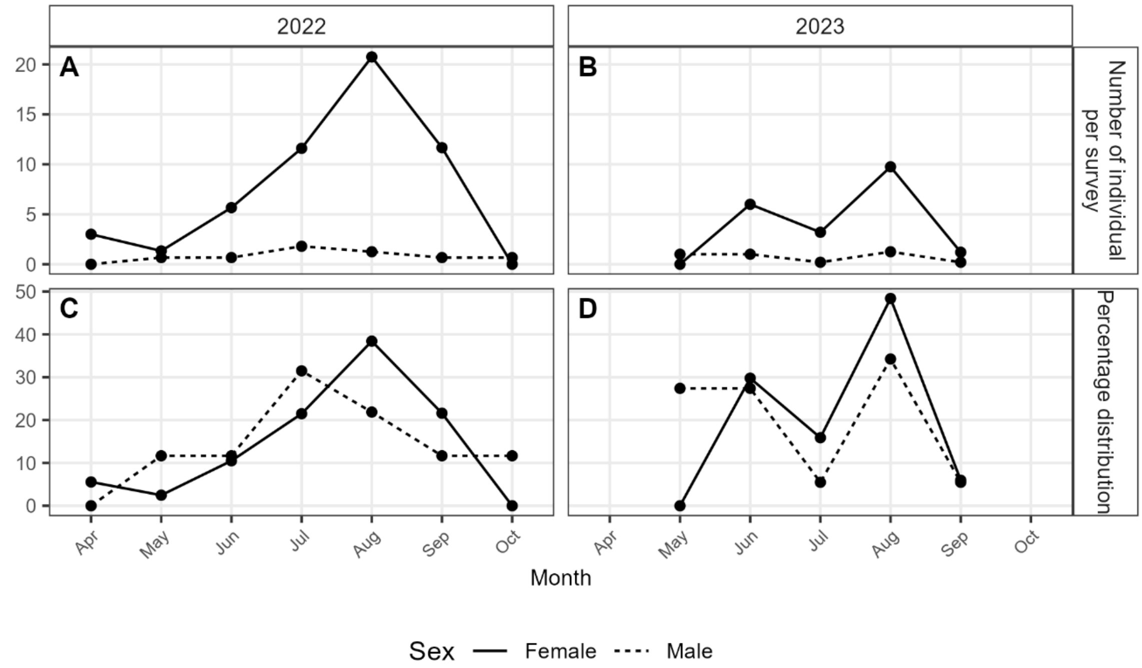

**Supplementary Figure S5:**

**Outputs of population modelling using historical temperatures and randomised temperatures (null model).** (A) Adjusted $R^2$ values for relationships between modelled nest counts and historical temperatures (blue) or randomised temperatures (yellow). (B) Predicted sex-ratio trajectories under the historical-temperature and null models. The red vertical line indicates the end of the 250-year burn-in period.

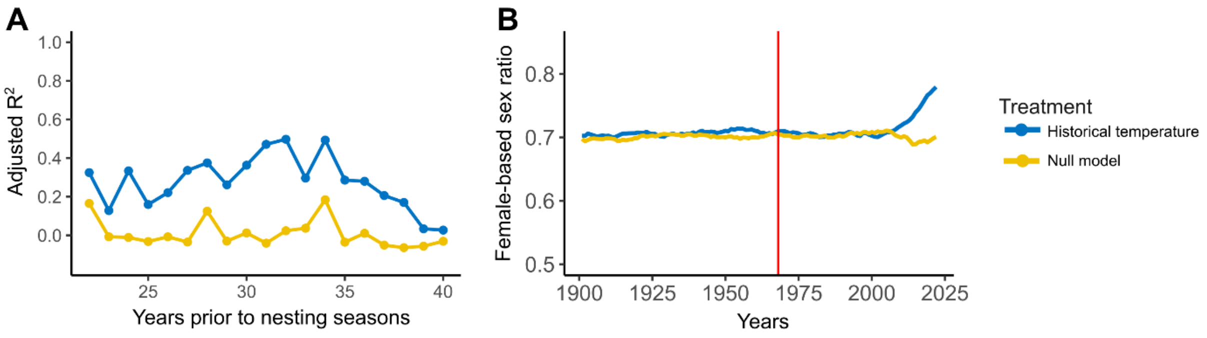

### Supplementary Figure S6:

**Linear relationships between log-transformed nest counts and historical air temperature at the best supported time-lag for each population**. Nest count data were obtained from the sources listed in Table 1 of the main manuscript. Temperature data were retrieved from the NOAA database described in the Methods. Both nest count and temperature datasets are available in the Zenodo repository (https://doi.org/10.5281/zenodo.24088945). USA: United States of America.

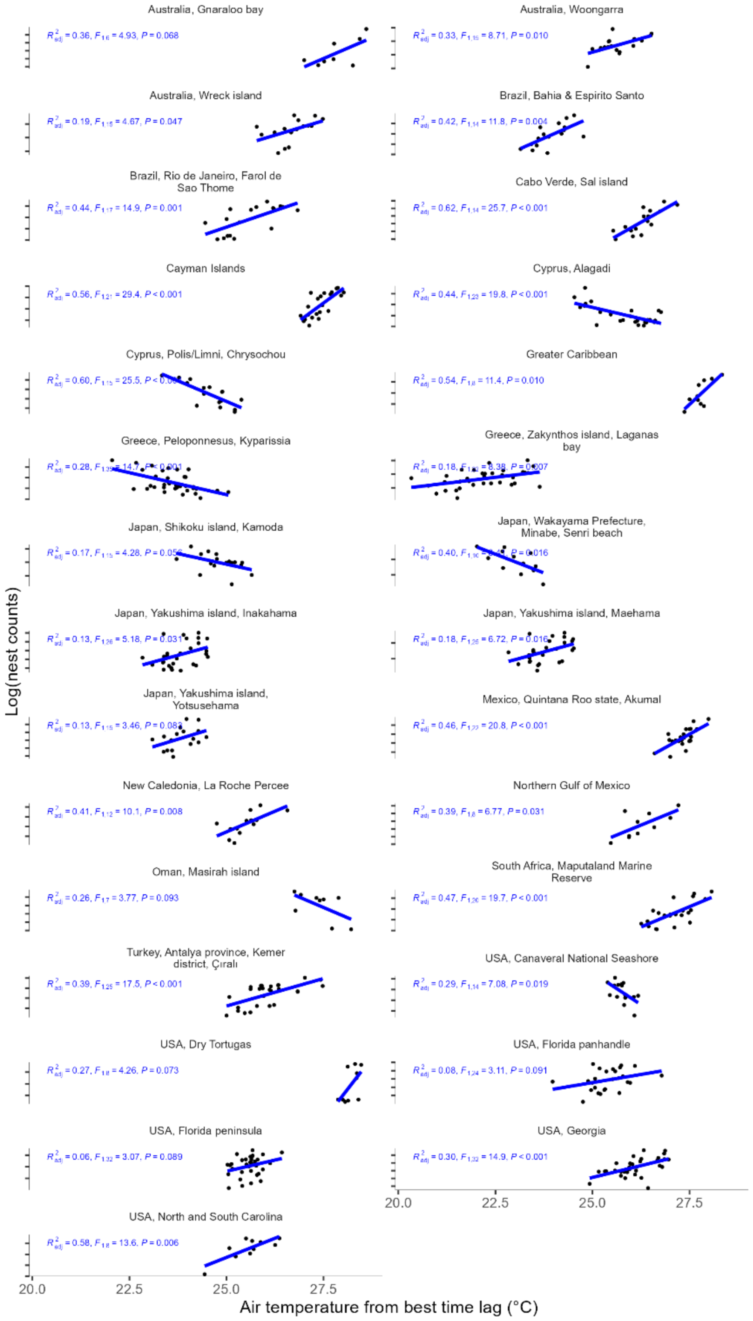

**Supplementary Figure S7:**

**Linear relationships between mean PTR and (A) latitude, (B) interannual temperature variability around the trend, and (C) the temperature trend**. The temperature trend was quantified as the slope of a linear regression of temperature against year, while interannual temperature variability was calculated as the standard deviation of the residuals around the fitted trend.

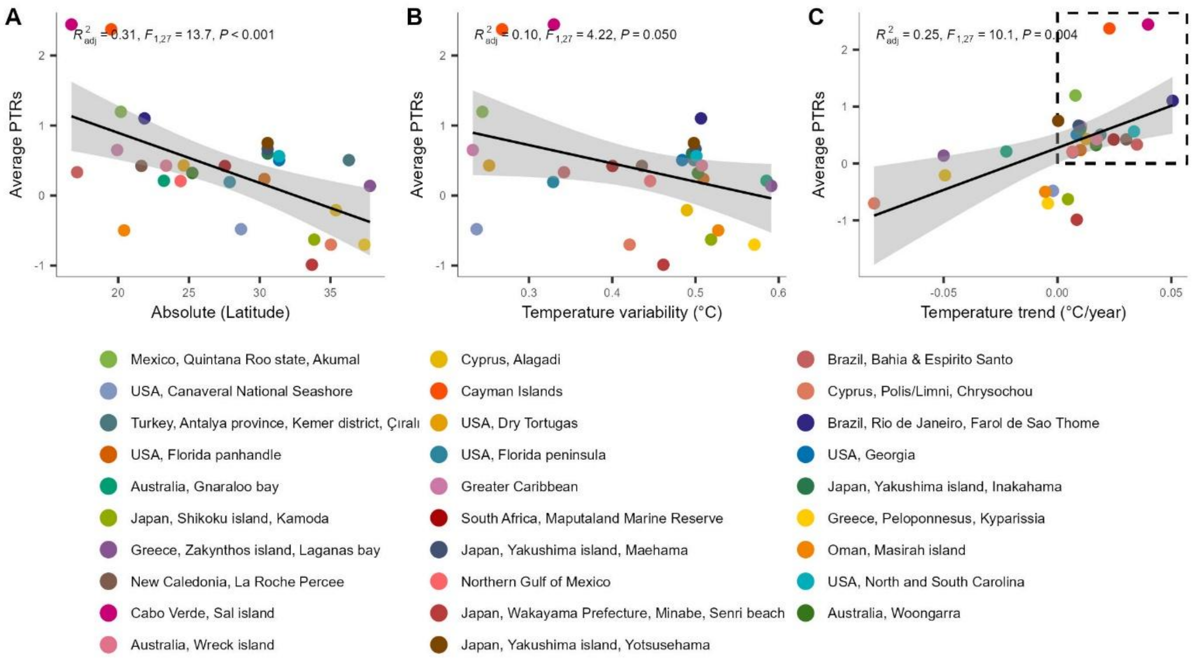
